# Mg^2+^ modulates the activity of hyperpolarization-activated calcium currents in plant cells

**DOI:** 10.1101/2020.01.14.906123

**Authors:** Fouad Lemtiri-Chlieh, Stefan T. Arold, Chris Gehring

## Abstract

Hyperpolarization-activated calcium channels (HACCs) are found in the plasma membrane and tonoplast of many plant cell types where they have an important role in Ca^2+^-dependent signaling. The unusual gating properties of HACCs in plants, i.e., activation by membrane hyperpolarization rather than depolarization, dictates that HACCs are normally open at physiological hyperpolarized resting membrane potentials (the so called pump or P-state), thus, if not regulated, they would be continuously leaking Ca^2+^ into cells. In guard cells, HACCs are permeable to Ca^2+^, Ba^2+^ and Mg^2+^, activated by H_2_O_2_ and the plant hormone abscisic acid (ABA) and their activity is greatly reduced by low amounts of free cytosolic Ca^2+^ ([Ca^2+^]_Cyt_) and hence will close during [Ca^2+^]_Cyt_ surges. Here we demonstrate that the presence of the commonly used Mg-ATP inside the cell greatly reduces HACC activity especially at voltages ≤ −200 mV and that Mg^2+^ causes this block. We therefore conclude, firstly, that physiological cytosolic Mg^2+^ levels affect HACCs gating and that channel opening requires either high negative voltages (≥ −200 mV) and/or displacement of Mg^2+^ away from the immediate vicinity of the channel. Secondly, based on structural comparisons with Mg^2+^-sensitive animal inward-rectifying K^+^ channel, we propose that the likely candidate HACCS described here are cyclic nucleotide gated channels (CNGCs), many of which also contain a conserved di-acidic Mg^2+^-binding motif within their pores. This conclusion is consistent with the electrophysiological data. Finally, we propose that Mg^2+^, much like in animal cells, is an important component in Ca^2+^ signalling and homeostasis in plants.

## INTRODUCTION

Ca^2+^ has long been recognized as an essential component in many plant cellular processes and in order for Ca^2+^ to function as an intracellular signal, temporal, spatial and stimulus specific changes in [Ca^2+^]_Cyt_ need to be tightly controlled (1). Ca^2+^-influx into plant cells is achieved, at least in parts, by three main types of Ca^2+^-channels (2,3): Firstly, channels that show little or no voltage sensitivity, referred to as non-selective calcium channels (NSCCs) primarily for being active at all voltages and for being less selective for calcium over monovalent cations such as K^+^ and Na^+^ (4); secondly, channels that show high voltage-dependence like those activated by depolarization (DACCs) (5) and thirdly others, which are the focus of this report, activated only by membrane hyperpolarization (HACCs) (6).

In guard cells, Ca^2+^ was shown to be involved in abscisic acid (ABA) signaling and stomatal guard cell movements (7,8) and using the patch clamp technique in both whole cell (WC) and excised configurations (EC), two types of HACCs were identified at the plasma membrane (PM). One type of HACCs is highly selective for Ca^2+^ (over K^+^ and Cl^−^) (9). Its activity is enhanced by ABA (9,10), H_2_O_2_ (10), and by external Ca^2+^ itself, an apparently unique property of this particular plant HACC (11). Meanwhile, increasing [Ca^2+^]_Cyt_ from 0.2 to 2 μM decreased the open probability (*P_o_*) of this HACC by a factor of 10 (9), implying a critical role of [Ca^2+^]_Cyt_ in the feedback regulation of the channel. The proteins responsible for this type of channel have not being identified to-date. This is not the case for the other type of HACCs where a family of 20 genes is known and their translation products were originally described as being mostly gated open by cyclic nucleotides such as 3’,5’-cyclic adenosine monophosphate (cAMP) or 3’,5’-cyclic guanosine monophate (cGMP) (12), hence their name: cyclic nucleotide gated channels (or CNGCs). Moreover, these plant channels, like their animal functional homologs, poorly discriminate between divalent and monovalent cations. Indeed, when heterologously expressed in oocytes, these channels are not only permeable to Ca^2+^ (13) but found to be equally permeable to monovalent cations such as K^+^, Rb^+^, Na^+^, Li^+^ and Cs^+^ (13,14). Similar inwardly-rectifying currents permeable to either Ca^2+^, Ba^2+^ and even Na^+^, activated by cAMP (15) and cGMP (16) were also characterized in guard cells with the patch clamp technique. The latter work demonstrated that the highly expressed CNGC5 and CNGC6 genes in guard cells are directly responsible for the recorded HACC activity. Likewise, in pollen tubes of Asian pear tree (*Pyrus pyrifolia* Nakai cv. Hosui), a HACC that conducts indiscriminately either Ca^2+^ or K^+^ was shown to be activated by cAMP and down regulated by high [Ca^2+^]_Cyt_ (17). It is noteworthy that all these type-two HACCs share one common characteristic, they are sensitive to low concentrations of Lanthanides (10,15-17).

We know from previous work, particularly in animal systems, that Mg^2+^ ions are key regulators of many ion channels and receptors (18). For instance, the inwardly rectifying K^+^ (Kir) channels and TRPV6, a member belonging to a subgroup of transient receptor potential (TRP) cation channels, both show strong Mg^2+^-dependent gating (19,20). Another example of a voltage-dependent block by Mg^2+^ ions is the N-methyl-D-aspartate (NMDA) receptor (21,22); in this case extracellular Mg^2+^ is responsible for this effect. By contrast, in plants, the rectification of *I*_K,in_ was found not to be due to a Mg^2+^-dependent block (23) but rather to an intrinsic property of the channel protein itself (24). However, ion channels localized to the tonoplast with a key role in stomatal volume regulation, such as the slow- (SV) and the fast- (FV) activating vacuolar channels, were shown to be affected by cytosolic Mg^2+^ ions (25–28). Indeed, it was found that, besides Ca^2+^, Mg^2+^ also promoted the activation of SV channels affecting their kinetics (time constants of channel activation and de-activation) and voltage-dependent activation characteristics. At the same time, Mg^2+^ inhibits FV channels, thus reducing K^+^ leakage from the tonoplast (25). Mg^2+^ was also shown to inhibit an outward NSCC, termed MgC, which was characterized in the PM of guard and subsidiary cells of *Vicia faba* and *Zea mays* (29). Another NSCC example from N_2_-fixing plants where Mg^2+^ plays a critical role is related to the transport of ammonium (NH_4_^+^)/ammonia (NH_3_) across the peribacteroid membrane (30). This is achieved in part through a passive non-selective electrogenic transport system, regulated by Ca^2+^, but more recently, this channel was described as having ‘unusual’ characteristics such as an inward rectification caused by Mg^2+^ on the cytosolic face and a very low single channel conductance (< 0.2 pS with 150 mM KCl + 10 mM CaCl_2_ in the pipette and 150 mM NH_4_^+^ in the bath) which was found to be inhibited by Mg^2+^ from the luminal face of the symbiosome (31).

ATP is another major regulator of ion channel gating. One of the best characterized channels is the K_ATP_, an inward K^+^-rectifier (from the K_ir_ family of genes) found mostly in cardiac and skeletal muscles, neurons and pancreatic β-cells (18). These channels are normally closed in the presence of ATP and only open, to hyperpolarize the cell membrane, when cytosolic ATP levels drop. ADP added in the form of Mg-ADP can restore the activity of K_ATP_ that were pre-treated with ATP (32,33). In guard cells, Mg-ATP is required for blue light-activated outward currents (34). Indeed, it was found that 1 to 2 mM Mg-ATP, as well as other intracellular substrates, are required to fully activate a plasma membrane electrogenic ion pump capable of hyperpolarizing the membrane to around −140 mV, a potential well beyond the activation threshold for *I*_K,in_ (35).

Given the importance and the key role that calcium channels play in plant cellular signaling including guard cell aperture regulation, we address the question of whether internal Mg^2+^ can affect HACCs activity in guard cell protoplasts (GCPs) and show examples of the Gd^3+^-sensitive Ba^2+^-currents (*I*_Ba_) activated by hyperpolarization with and without Mg^2+^ in the patch pipet. We also assess important properties of *I*_Ba_ in the absence of Mg^2+^ such as the permeability and sensitivity to some relevant inorganic compounds and physiological effectors such as abscisic acid (ABA) and cAMP.

## EXPERIMENTAL PROCEDURES

### Protoplast isolation

*Vicia faba* L. *cv* (Bunyan) Bunyan Exhibition seeds were grown on vermiculite under conditions described previously (36). *Arabidopsis thaliana* (Columbia) seeds were grown on peat pellets (jiffy, Oslo) in a controlled environment growth chamber (Percival, CLF plant climatic, Wertingen) at 22 °C on a 8/16-h light/dark cycle. Guard cell protoplasts (GCPs) were isolated from either 3- to 4- week *V. faba* or 5- to 6- week *A. thaliana* plants. GCPs were isolated from abaxial epidermal strips as described previously (37). Briefly, epidermal strips were floated on medium containing 1.8-2.5% (w/v) Cellulase Onozuka RS (Yacult Honsha, Tokyo, Japan), 1.7-2% (w/v) Cellulysin (Calbiochem, Behring Diagnostics, La Jolla, CA), 0.026 % (w/v) Pectolyase Y-23, 0.26 % (w/v) BSA, and 1 mM CaCl_2_ (pH 5.6) with osmolality adjusted with sorbitol to 360 mOsm.kg^−1^. After 90-120 min incubation in the dark at 28 °C with gentle shaking, released protoplasts were passed through a 30 μm mesh, and kept on ice for 2 to 3 min before centrifugation (100 g for 4 min at room temperature). The pellet consisting of GCPs was re-suspended and kept on ice in 1 or 2 ml of fresh medium containing 0.42 M mannitol, 10 mM 2-(N-morpholino)ethanesulfonic acid (Mes), 200 μM CaCl_2_, 2.5 mM KOH (pH 5.55 and osmolality at 466 mOsm.kg^−1^). Unless stated otherwise, all chemicals were from Sigma (Sigma-Aldrich Co. St Louis, MO).

### Solutions

Protoplasts were placed in a 0.5 ml chamber, left to settle down, and then perfused continuously at flow rate of ≈ 0.5-1 ml/min. To record *I_Ba_* currents through HACC, we used barium containing solutions. The bath medium contained (in mM): 100 BaCl_2_, 10 Mes (pH 5.5 with Tris base) and the pipette contained (in mM): 100 BaCl_2_, 4 EGTA, 10 Hepes (pH 7.5 with Tris). Experiments where *I*_K,in_ and *I*_Ba_ measurements were made on the same GCPs a different bath and internal solutions as follows. Bath (in mM): 30 KCl, 10 Mes (pH 5.5 with Tris base) to measure *I*_K,in_ and was replaced by 100 BaCl_2_, 10 Mes (pH 5.5 with Tris base) to measure *I*_Ba_. Internal solution (in mM): 1 BaCl_2_, 18 KCl, 4 EGTA, 10 Hepes (pH 7.5 with Tris base). Mg-ATP, MgCl_2_ and/or K_2_-ATP were added as specified in the figures. Osmolality was adjusted with Sorbitol to 210mOsm.kg^−1^. For classic solutions used to measure *I*_K,in_ (Fig. 4 A, and B), refer to (38). ABA was added externally. All chemicals were from Sigma Chemical, Poole Dorset, UK. The membrane permeable cAMP analog Bt_2_cAMP was solubilized in deionized water and stored in aliquots of 50–100 μl at a concentration of 0.1 M. Bt2cAMP was diluted to the final desired concentration just a few minutes before its use.

### Current-voltage recording and analysis

Patch pipettes (5-10) were pulled from Kimax-51 glass capillaries (Kimble 34500; Kimble, Owens-Illinois) using a two stage puller (Narishige PP-83, Japan). Experiments were performed at room temperature (20 to 22 ^o^C) using standard whole-cell patch clamp techniques, with an Axopatch 200B Integrating Patch Clamp amplifier (Axon Instruments, Inc. Union City, CA, U.S.A.). Voltage commands and simultaneous signal recordings and analyses were assessed by a microcomputer connected to the amplifier via multipurpose Input/output device (Digidata 1320A) using pClamp software (versions 8.0 and 10; Axon Instruments, Inc.). After Giga-Ohm seals are formed, the whole-cell configuration was then achieved by gentle suction, and the membrane was immediately clamped to a holding voltage (*h*_v_) of −36 mV. GCPs were continuously perfused throughout the experiment and current recordings began only after at least 5-10 min from going into whole-cell mode to allow for intracellular equilibrium between cytoplasm and patch pipet solution. All current traces shown were low-pass filtered at 2 kHz before analog-to-digital conversion and were uncorrected for leakage current or capacitive transients. Membrane potentials were corrected for liquid junction potential as described (39). Ionic activities were calculated using GEOCHEM-EZ (40). Current-voltage (I-V) relationships for *I*_Ba_ and *I*_K,in_ were plotted as steady-state currents *vs.* test potentials when using the square pulse stimulations or utilizing the “Trace *vs* Trace” feature of Clampfit analysis when using voltage ramps. Unless otherwise stated, every experiment reported here was repeated a minimum of three times and data were graphed as mean±s.e.m.

### Homology modeling

Models were built using Swiss Model and structures visualized with PyMOL (The PyMOL Molecular Graphics System, Version 1.5.0.4 Schrödinger, LLC). Multi-sequence alignments were produced with Muscle (43).

## RESULTS

### Removal of intracellular Mg-ATP unveils a larger instantaneously-activated, inwardly-directed and Gd^3+^- sensitive Ba^2+^-current

We recorded Ba^2+^-currents (*I*_Ba_) in guard cell protoplasts either in the presence or absence of Mg-ATP (Fig. 1A) and at the end of the trials we also tested for *I*_Ba_ sensitivity to Gd^3+^, a potent blocker of *I*_Ba_ (Fig. 1B). The recorded currents were generated in response to a square pulse protocol from +64 mV to −256 mV in increments of −20 mV with the holding voltage (*h*_V_) set to −36 mV. In the presence of Mg-ATP (Fig.1 A; left panel), a small instantaneous inward-rectifying current of ≈ −20 pA started to activate around −200 mV to only reach a maximum of −60 pA at −256 mV. This current was, as expected, sensitive to the addition of extracellular Gd^3+^ (see Fig. 1B; left panel). Meanwhile in the absence of Mg-ATP (Fig. 1A; right panel), much larger (≥ 8-fold) instantaneous rectifying-currents, which happened to be also Gd^3+^-sensitive were recorded (Fig. 1B; left traces). The current-voltage (I-V) relationships in the presence (●; n=3) or absence (○; n =7) of Mg-ATP are represented in Figure 1C (left panel). We also plotted the effect of Gd^3+^ on the I-V relationships in the presence (■) or absence (□) of Mg-ATP (Fig. 1C; right panel). These results highlight that when omitting Mg-ATP from the intracellular medium, a larger Gd^3+^- sensitive inwardly-rectifying Ba^2+^-current is unveiled that activates at significantly less negative voltages (see the shift to the right of ≥ −100 mV in the I-V plot). Furthermore, the currents recorded in 0 Mg-ATP seem to reverse near the calculated Nernst equilibrium potential for Ba^2+^ (*E*_Ba_ ≈ +28 mV), and are far removed from *E*_Cl_ (−54 mV). Likewise, using fast depolarization ramps (0.07 V/sec) after activating the current *I*_Ba_ with a square pulse to −156 mV (see voltage protocol and current trajectories in Fig. 1D; left), a reversal potential of +17 mV was measured, again close to *E*_Ba_ rather than *E*_Cl_ (see Fig. 1D, zoomed I-V plot). This is also indicative of the higher permeability of this conductance to Ba^2+^ as compared to Cl^−^.

**Figure 1.**
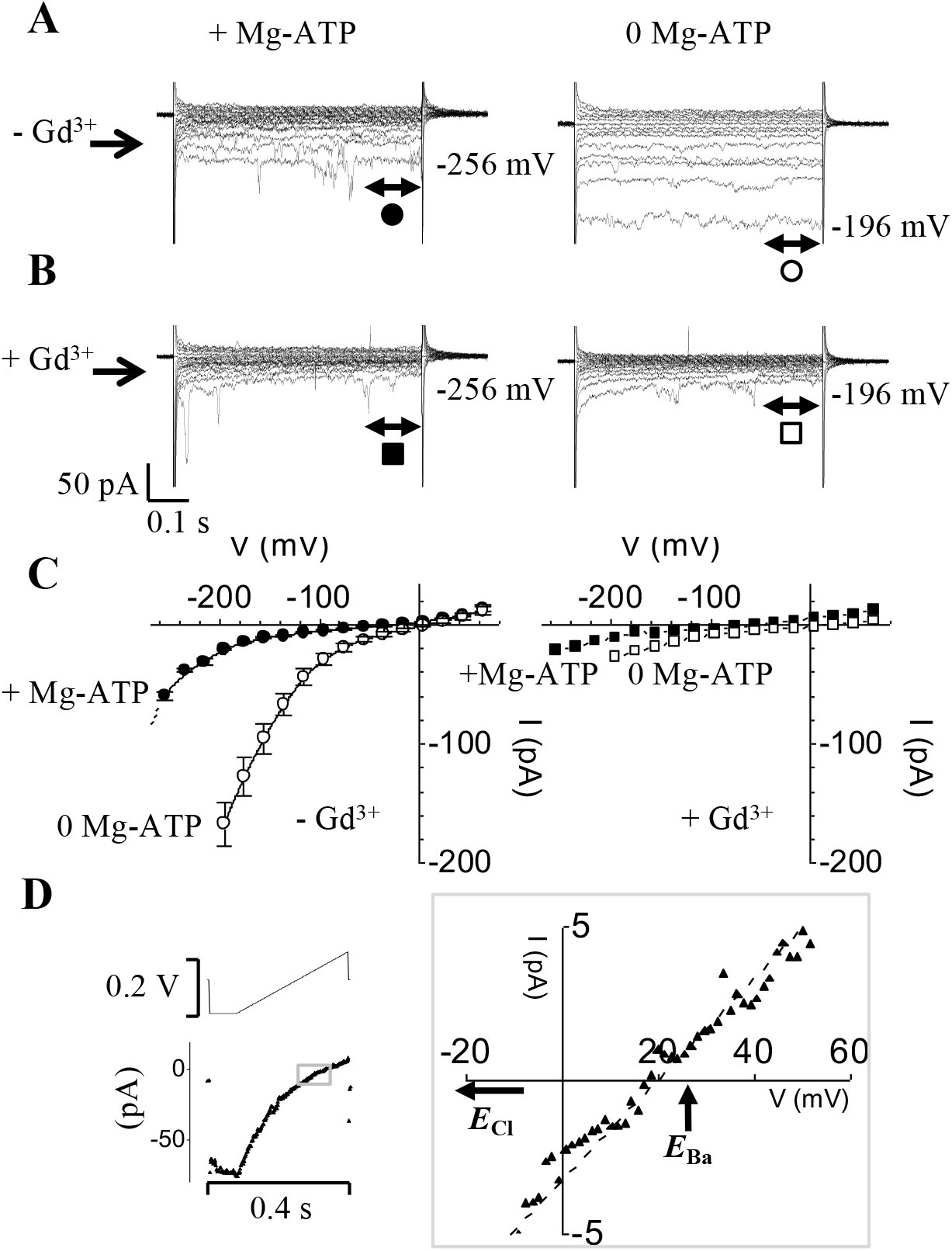
Removal of intracellular Mg-ATP unveils a larger instantaneously-activated, inwardly-directed and Gd^3+^-sensitive Ba^2+^ current. (**A**) Typical examples of current traces in whole-cell mode obtained from two separate guard cells with the pipette solution either containing (*left traces*) or lacking (*right traces*) Mg-ATP (1 mM). The pulse protocol mostly used throughout this study consists of 0.6 s long square voltage pulses ranging from +64 to −256 mV in −20 mV increments; the holding potential *h*_v_ was set to −36 mV. In order to preserve the quality of the ‘Giga’ seals, GCPs with no Mg-ATP in the pipette were not subjected to higher voltages beyond −196 mV. (**B**) Current traces of *I*_Ba_ obtained from the same cells and in the same conditions as described in (A) except for the external solution containing Gd^3+^ (20 μM; left or 100 μM; right) (**C**) Current-voltage relationships (I-V); *left panel*: superimposed I-Vs of *I*_Ba_ in the absence (○; n=7) or presence (●: n=3) of Mg-ATP. *Right panel*: I-V relations in the presence of Gd^3+^ (■: +Mg-ATP; □: -Mg-ATP). (**D**) Typical current trace recorded in the absence of Mg-ATP (below) in response to the ‘*E*_rev_ protocol’ which consists of activating *I*_Ba_ using a hyperpolarization square pulse to −156 mV and immediately followed by a continuous depolarizing ramp to +64 mV (the voltage protocol is depicted above) with a slope of 0.7 V s^−1^. Scale bars are shown below the current traces and to the left of the voltage protocol.

To test whether changing external Ba^2+^ concentration will affect current magnitude as well as the I-V relationship, GCPs were patched in whole cell mode using the Mg-ATP free internal solution (Fig. 2). Once again, the current magnitudes recorded in the absence of Mg-ATP are quite substantial. For instance, at −196 mV, a −280 pA current is measured in 100 mM [Ba^2+^]_out_ (Fig. 2A) while in 30 mM [Ba^2+^]_out_ (Fig. 2B) the same voltage gives rise to a current value of −220 pA. All current magnitudes at any given voltage are decreased when switching to lower [Ba^2+^] in the bath. The corresponding I-V plots appear as shifting to negative values (Fig. 2C) when switching from 100 to 30 mM Ba^2+^ and this is accompanied by a negative shift in the apparent reversal potential (*E*_rev_) values. Indeed, when zooming in (Fig. 2C, inset), the apparent *E*_rev_ shows a negative shift of ≈ −14 mV as a result of this Ba^2+^ concentration change. Furthermore, the apparent *E*_rev_ in 30 mM [Ba^2+^]_out_ (≈ +8 mV) is still closer to the calculated *E*_Ba_ rather than *E*_Cl_ which are in this case +17.7 mV and −25.7 mV respectively. This experiment was repeated with 100, 30 and 10 mM BaCl_2_ in the bath and the same qualitative effects were seen i.e., decrease of current amplitude when decreasing the [Ba2+]_o_ and negative shift in the apparent *E*_rev_ (data not shown).

### HACC permeability sequence to divalent cations in the absence of Mg-ATP: Ba > Ca ≈ Sr ≈ Mn ≫ Mg

Guard cell permeability to other divalent cations such as Ca^2+^, Sr^2+^, Mn^2+^ and Mg^2+^ in the absence of Mg-ATP was also tested (Fig. 3). As expected, HACC was permeable to Ca^2+^ (Fig. 3A) and found to have similar permeability to both Sr^2+^ (Fig. 3A) and Mn^2+^ (Fig. 3B). Meanwhile Mg^2+^ did not permeate HACC (Fig. 3A). The I-V plots (Fig. 3A and B) summarizes the permeability data: i.e., the lack of HACC permeability to Mg^2+^ as well as the much larger permeability to Ba^2+^ when compared to either Ca^2+^, Sr^2+^ or Mn^2+^. The effect of Mn^2+^ ions over time is reported in Figure 3B and highlights the unique behavior of this ion. Unlike Ca^2+^ or Sr^2+^, Mn^2+^ (Fig. 3B) triggered a transient block of HACC followed by some current recovery while still washing out the Ba^2+^ and replacing it with 100 mM Mn^2+^. This transient block effect was repeated on two other GCPs but was never seen with either Ca^2+^ or Sr^2+^; nor was it seen with Mg^2+^ even after 20 minutes of washing out Ba^2+^.

**Figure 2.**
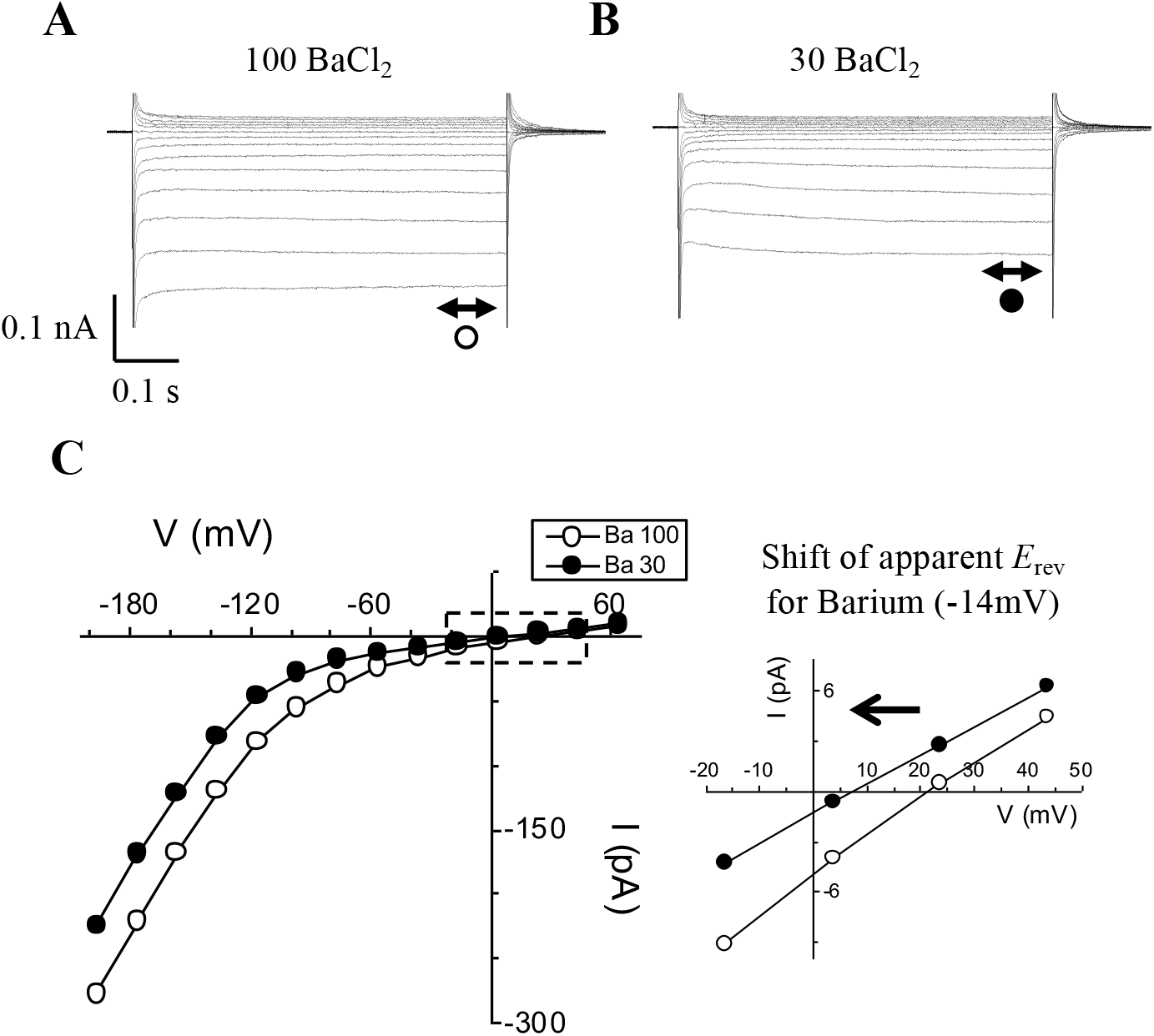
Decreasing [Ba^2+^]_o_ from 100 to 30 mM not only decreases the hyperpolarization-activated *I*_Ba_ but also shifts its apparent *E*_rev_ (~-14 mV). (**A**) and (**B**) Typical current traces of hyperpolarization-activated *I*_Ba_ recorded from the same guard cell either in 100 (A) or 30 mM (B) [Ba^2+^]_o_. Scale bars are shown below the current traces. (**C**) Corresponding I-V plots of *I*_Ba_ in 100 and 30 mM [Ba^2+^]_o_ taken from the current traces shown in (A) and (B). Inset: Zoomed I-V plot from the the area shown as a dashed box in (C). The inset shows the amount (in mV) and the direction (arrow) of the shift in the apparent *E*_rev_ when the bath perfusion was switched from 100 to 30 mM Ba^2+^.

**Figure 3.**
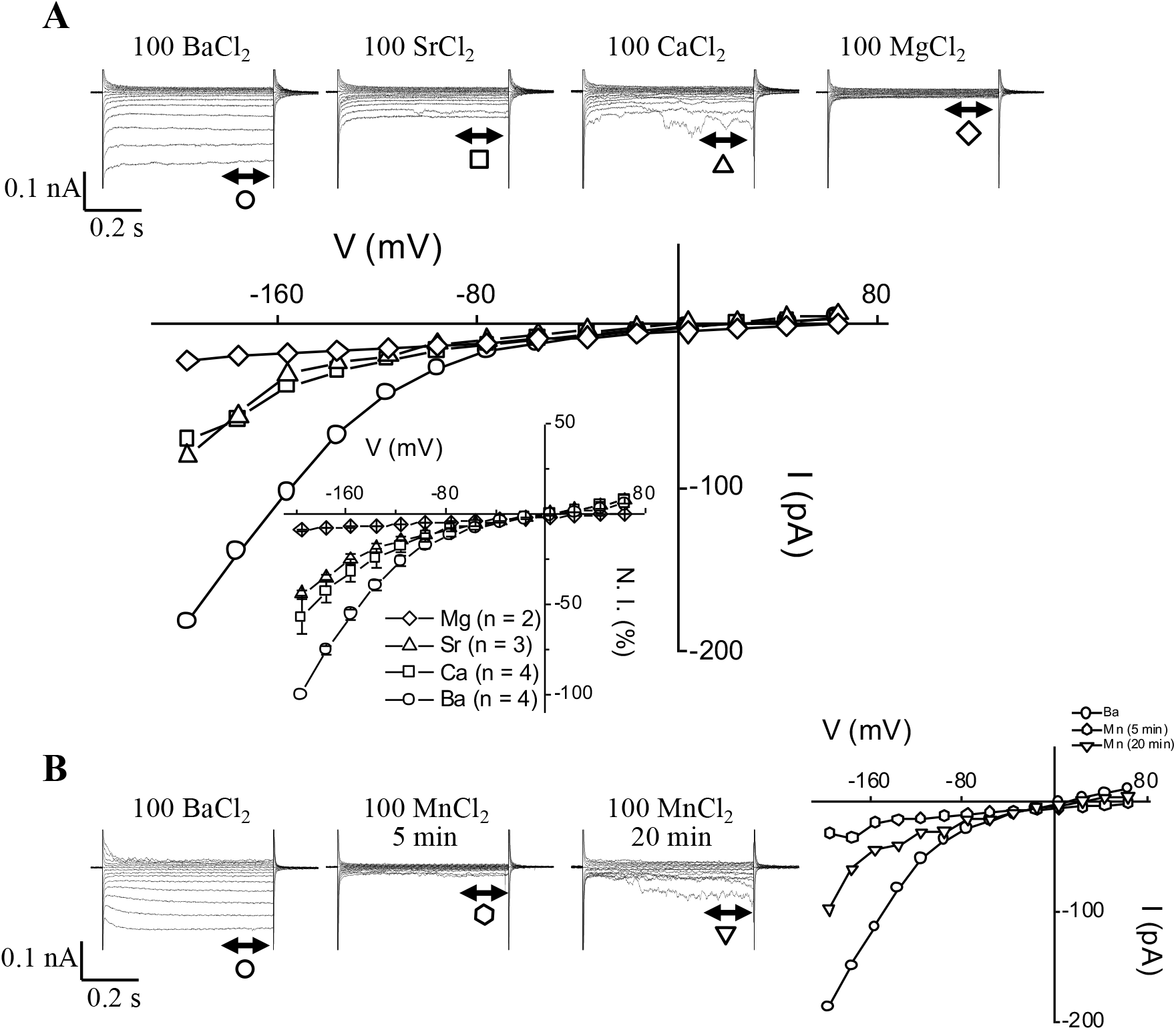
Current through *I*_Ba_ channels can be carried by other divalent cations such as Ca^2+^, Sr^2+^ and even Mn^2+^ but not by Mg^2+^. Typical current traces and corresponding I-V plots recorded in the presence of 100 mM BaCl_2_, 100 mM CaCl_2_, 100 mM SrCl_2_ and 100 mM MgCl_2_ (note that all traces are from the same guard cell except for MgCl_2_). Inset: Normalized group I-V curves showing divalent permeabilities (the mean current values obtained for Ca^2+^, Sr^2+^ and Mg^2+^ were normalized to the mean current value obtained in Ba^2+^ at −196 mV). (B) Typical current traces and corresponding I-V plots recorded in the presence of 100 mM BaCl_2_ and 100 MnCl_2_ at times 5 and 20 minutes. All traces are from the same guard cell. Notice the transient blocking effect of Mn^2+^ ions.

**Figure 4.**
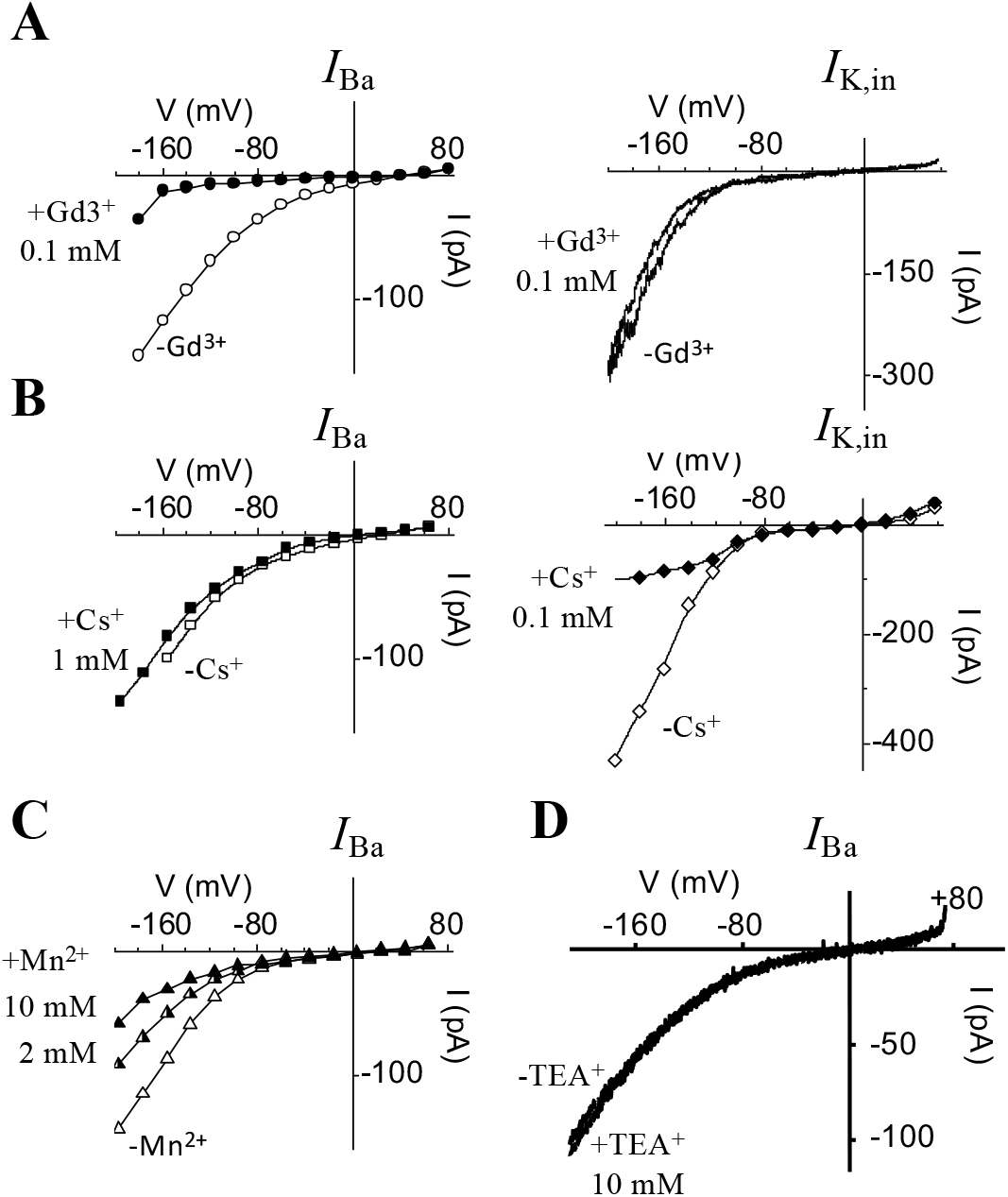
Differential effects of some known blockers on the two main PM conductances activated by hyperpolarization in guard cells: namely, *I*_Ba_ and *I*_K,in_. (**A**) I-V plots showing the effect of Gd^3+^ (0.1 mM) on *I*_Ba_ (*left panel*) and *I*_K,in_ (*right panel*). (**B**) I-V plots showing the effect of 1 mM Cs^+^ on *I*_Ba_ (*left panel*) compared to the effect of 0.1 mM Cs^+^ on *I*_K,in_ (*right panel*). (**C**) I-V plot showing the effect of Mn^2+^ on *I*_Ba_ recorded from the same GCP using 2 and 10 mM in the bath. (**D**) *I*_Ba_-V plots generated from current recordings using hyperpolarizing ramps (+64 to −196 mV; 0.7 V.s^−1^; *h*_v_ = −36 mV) showing the effect of 10 mM TEA^+^ added to the bath.

### HACC permeability sequence to monovalent cations in the absence of Mg-ATP: K≈Na>Ba>Cs≫TEA

We observed that this Gd^3+^-sensitive HACC is also permeable to some physiologically relevant monovalent cations such K^+^ (Suppl. Fig. 1A), Na^+^ (Suppl. Fig. 1B) and Cs^+^ (Suppl. Fig. 1C) but not tetraethylammonium (TEA^+^)(Suppl. Fig. 1D). These data indicate that the Gd^3+^-sensitive current characterized in this work does not select for small mono or divalent cations (except for the case of Mg^2+^ and the bigger cation TEA^+^). Chloride too does not seem to permeate through this HACC. Indeed, when 100 mM Cl^−^ was added at the same time as TEA^+^ (see Suppl. Fig. 1D), no current could be detected, indicating that Cl^−^ is as impermeable as TEA^+^. Qualitatively, the same effect was seen in all patched GCPs (n = 5 for K^+^ and Na^+^; n = 2 for Cs^+^ and TEA^+^).

### Effect of blockers of I_Ba_ in the absence of Mg-ATP and comparison with the effect on I_K,in_

In order to further characterize this HACC, readily unmasked when Mg-ATP was omitted from the patch pipet, the effect of some classical blockers such as the lanthanides (La^3+^ and Gd^3+^), Mn^2+^, Cs^+^ and TEA^+^ were tested on *I*_Ba_ as well as on *I*_K,in,_ (the other major conductance that activates upon hyperpolarization in guard cells). One of the most conspicuous effects lies in the potent effect of Gd^3+^ in blocking *I*_Ba_ (Fig. 4A) even when used at relatively low concentrations (20 to 100 μM) while the same concentrations of Gd^3+^ had no effect on *I*_K,in_ (Fig. 4A). An even higher concentration of Gd^3+^ (500 μM) did not affect *I*_K,in_ (data not shown). La^3+^ also blocked *I*_Ba_ measured in 0 Mg-ATP, but we found that much higher concentrations of La^3+^ (0.2 to 0.5 mM) are needed to achieve the same block as compared to Gd^3+^ (data not shown). Experimenting with cesium, a blocker of *I*_K,in,_ hardly any effect on *I*_Ba_ was registered (Fig. 4B). Even though Cs^+^ was used at concentrations up to 1 mM, it had only a small effect, if any, on *I*_Ba_ while the tenth of this amount (0.1 mM) is sufficient to block a large proportion (≈ 80% or more) of *I*_K,in_ (Fig. 4B). Furthermore, Mn^2+^ used at 2 mM inhibited HACC by ≈ 37% (at V = −196 mV) when the charge carrier (in this case Ba^2+^) was still present in the bath (Fig. 4C). Increasing Mn^2+^ concentration to 10 mM shows that Mn^2+^ is not an efficient blocker of *I*_Ba_ as compared to Gd^3+^ or La^3+^ and 10 mM Mn^2+^ only causes an extra 20% *I*_Ba_ inhibition (see IV plot in Fig. 4C). Finally, 10 mM tetraethylammonium chloride (TEA), a concentration that was shown to block 70 to 80 % of *I*_K,in_ in intact guard cells (44), had no effect whatsoever on *I*_Ba_ (Fig. 4D) measured in 0 Mg-ATP.

### Rapid enhancement of I_Ba_ by ABA

Given that in guard cells, a HACC was implicated downstream of ABA in stomatal movements, we tested whether ABA affects *I*_Ba_ activated in the absence of internal Mg-ATP. We patch clamped guard cells to measure *I*_Ba_ currents under baseline conditions, i.e., zero Mg-ATP inside and no added ABA outside (Fig. 5). After about 10 minutes, the time usually necessary to reach steady-state conditions, we switched the perfusion solution to the one containing 20 μM ABA. A rapid and pronounced increase of *I*_Ba_ currents is seen at all voltages between −100 and −200 mV (≥ 1.3 to 1.5 fold)after only 5 minutes of ABA treatment (Fig. 5B) and a near doubling of the size of the *I*_Ba_ currents occurs at 10 minutes (Fig. 5, A and B). The enhancement of *I*_Ba_ in response to ABA, especially at 10 minutes, spans from −60 to −200 mV and also appears to shift the activation threshold of *I*_Ba_ (Fig. 5B) to the right. This suggests that ABA not only enhances calcium entry through HACC but can also mobilize calcium entry at less negative voltages.

**Figure 5.**
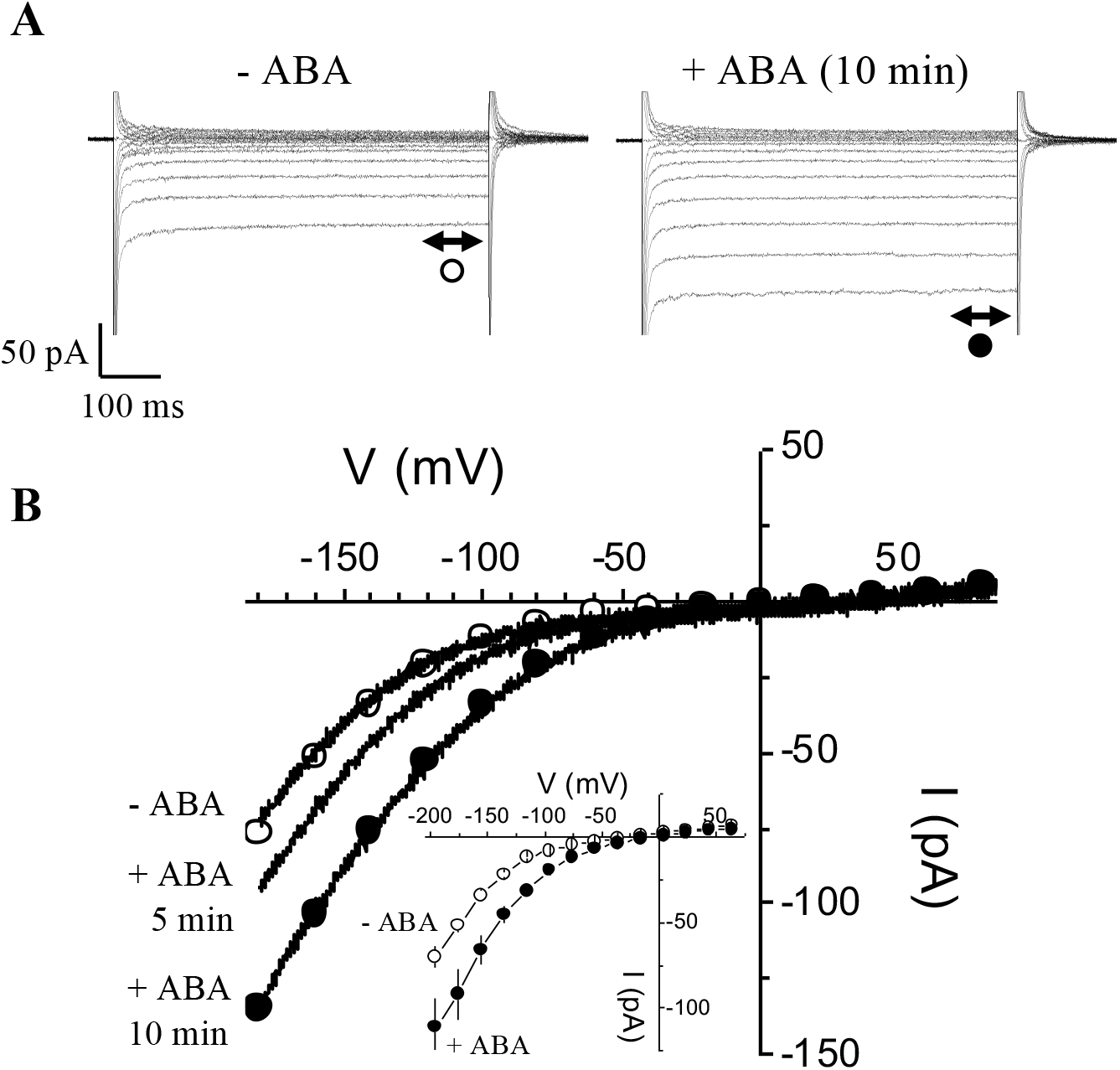
Rapid enhancement of *I*_Ba_ by ABA. (**A**)*I*_**Ba**_ currents in the absence of Mg-ATP in the patch pipet recorded from the same guard cell in response to hyperpolarizing voltages (from +64 to −196 mV; in −20 mV increments) before (○) and 10 minutes after (●) bath application of ABA (20 mM). (B) I-V plots of the effect of ABA showing the enhancing effect of ABA with time. (control: -ABA; 5 and 10 minutes after continuous bath perfusion with ABA). We also superimposed the measurements generated by voltage ramps for the control and 10 min ABA. Inset: Superimposed I-V plots showing the average effect of ABA on *I*_**Ba**_ (0 Mg-ATP). Data are current average measurements (± sem) from different experiments (n=3) before (○) and about 10 minutes after (●) bath perfusion with ABA (Student test; *: P≤0.05, **: P≤0.01, ns: P> 0.05).

### Characterization of the effect of Mg^2+^on I_Ba_ and I_Kin_

To answer whether internal Mg^2+^ alone causes the block of *I*_Ba_ when we add Mg-ATP, GCPs were patched either without Mg-ATP or without ATP but with added Mg^2+^ (as MgCl_2_). Furthermore, and as a control for ‘ion transport functionality’ of the patched GCPs, we used conditions that allow measurements of not just *I*_Ba_, but also to record *I*_K,in_. The experiments were started in conditions allowing to probe for *I*_K,in_ with KCl (30 mM) in the bath and this was then followed by replacing the KCl with a solution containing BaCl_2_ (100 mM). This was done first in the absence of both intracellular Mg^2+^ and ATP (Fig. 6A) and then repeated on another batch of guard cells with internal medium containing 1 mM MgCl_2_ but no ATP (Fig. 6B). Firstly, in zero Mg-ATP the only current that activated in response to hyperpolarization was *I*_Ba_ (Fig. 6A). *I*_K,in_ could not be activated. Secondly, when only Mg^2+^ was included in the pipet solution (no ATP added), *I*_K,in_ could now be activated in 30 mM K^+^, however, when switching the bath from K^+^- to Ba^2+^-containing media, *I*_Ba_ currents vanished (Fig. 6C) indicating that including only Mg^2+^ in the patch pipet can cause the block of *I*_Ba_ at voltages where they are normally activated in 0 Mg-ATP.

### Could cAMP activate HACC in guard cells despite the presence of intracellular Mg^2+^?

We know from our own previous work (15) that cAMP activated a Gd^3+^-sensitive HACC in guard cells while recording in Mg^2+^- and ATP-free media (see also, Suppl. Fig. 2A) but since we now show that this conductance discriminates poorly between divalent and monovalent cations, a hallmark characteristic of all animal as well as plant cyclic nucleotides gated channels (CNGCs) (13,45), we sought to check if the Gd^3+^-sensitive *I*_Ba_ is also gated by cAMP in conditions where intracellular Mg^2+^ is present and *I*_Ba_ is already blocked (Suppl. Fig. 2B). In the absence of db_2_cAMP (the lipophilic permeable analog of cAMP), and as expected only a small background *I*_Ba_ current is seen (≤10 pA around −190 mV). After perfusing with db_2_cAMP (1 mM), a substantial increase in *I*_Ba_ amplitude between voltages from around −30 to −190 mV was observed (>60 pA around −190 mV). Keeping db_2_cAMP in the bath and adding Gd^3+^ (50 μM) resulted in a total block of the current (≤4 pA around −190 mV). This may indicate that GCPs harbor CNGCs that can be activated by cAMP despite the blocking effect by Mg^2+^.

**Figure 6.**
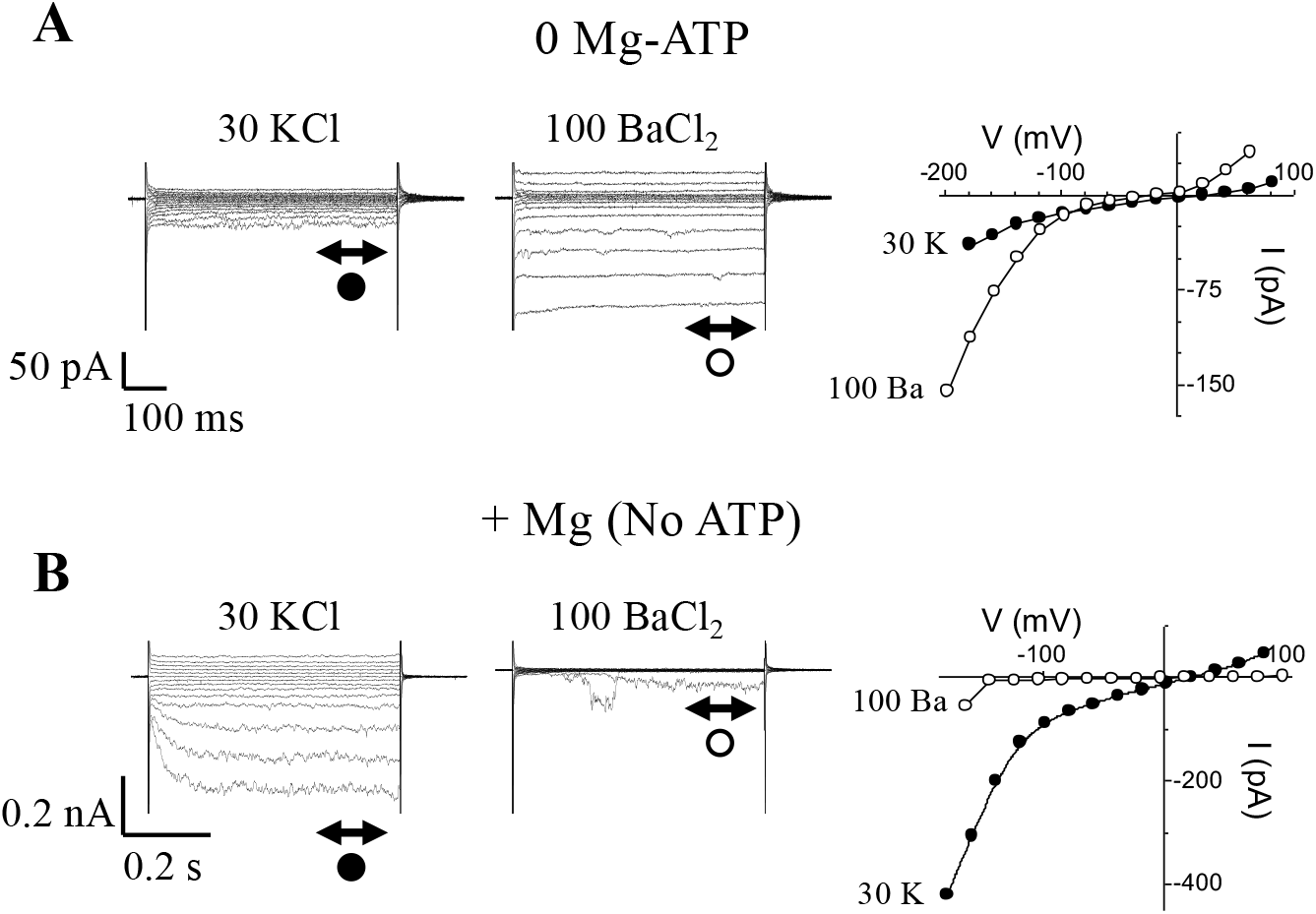
Internal Mg^2+^ is responsible for *I*_Ba_ inhibition. (**A**) Currents (*left panel*) and corresponding I-V plots (*right panel*) from the same guard cell recorded in response to hyperpolarizing voltage steps (from +64 to −196 mV; in −20 mV increments; *h*_v_ = −36 mV) and in the absence of Mg-ATP either using K^+^ (30 mM) or Ba^2+^ (100 mM) as charge carriers. Notice the larger *I*_Ba_ compared to *I*_K,in_. (**B**) Current (*left panel*) and corresponding I-V relationships (*right panel*) recorded in the same external conditions as in (A) but this time, using only Mg^2+^ ions in the patch solution (1 mM added as MgCl_2_) and no added ATP. In these conditions, the predominant current activated by hyperpolarization is *I*_K,in_, not *I*_Ba_.

### Identification of candidate Mg^2+^-dependent cation channels

Crystallographic and functional analyses of a strong inward-rectifier K^+^ channel (Kir2.2) from chicken (46) showed that the rectification characteristic can be explained by Mg^2+^-binding to negatively charged regions in the pore (formed by D173) with possible additional contributions from acidic residues within the cytoplasmic regulatory domains (D256 and E300/E225) (Suppl. Fig. 3). To explore the possibility that a similar mechanism allows Mg^2+^ inward rectification for a subset of candidate plant channels that display the electrophysiological properties described here (i.e. activation by hyperpolarization and cAMP, absence of cation selectivity and inhibition by lanthanides), we built homology models of the pore-forming residues for AtCNGCs (47). The models were built using structures of the human hyperpolarization-activated channels HCN1, based on a ~22 % sequence identity. The obtained models clearly showed that AtCNGCs do not have a Mg^2+^-binding site corresponding to the location of Kir2.2 D173 inside the pore region (Fig. 7A, left arrow, and Suppl. Fig. 3). However, a tandem glutamic acid motif that could form a possible di-acidic Mg^2+^-binding site is found in most AtCNGCs located a little downstream of the pore in the so-called C-linker disc (Fig. 7). Akin to Kir2.2 D173 and E300/E225, this di-acidic AtGNCG motif forms a positively charged opening with distances of 7.3–8.3Å between neighbouring charges and a diameter of ~11Å [for Kir2.2 D173 these values are 7.3–7.4Å and 10.4–11.8Å, respectively (PDB 3jyc)]. As in Kir2.2, these distances between carboxyl groups are too large for a direct ion coordination, suggesting that Mg^2+^ is bound through bridging water molecules (46). Interestingly, this di-acidic motif is not present in AtGNCG2, which has been shown to be an atypical family member with respect to ion selectivity (49). The di-acidic motif is also absent in HCN1, for which Mg^2+^ inward rectification has not been documented (Fig. 7C).

**Figure 7.**
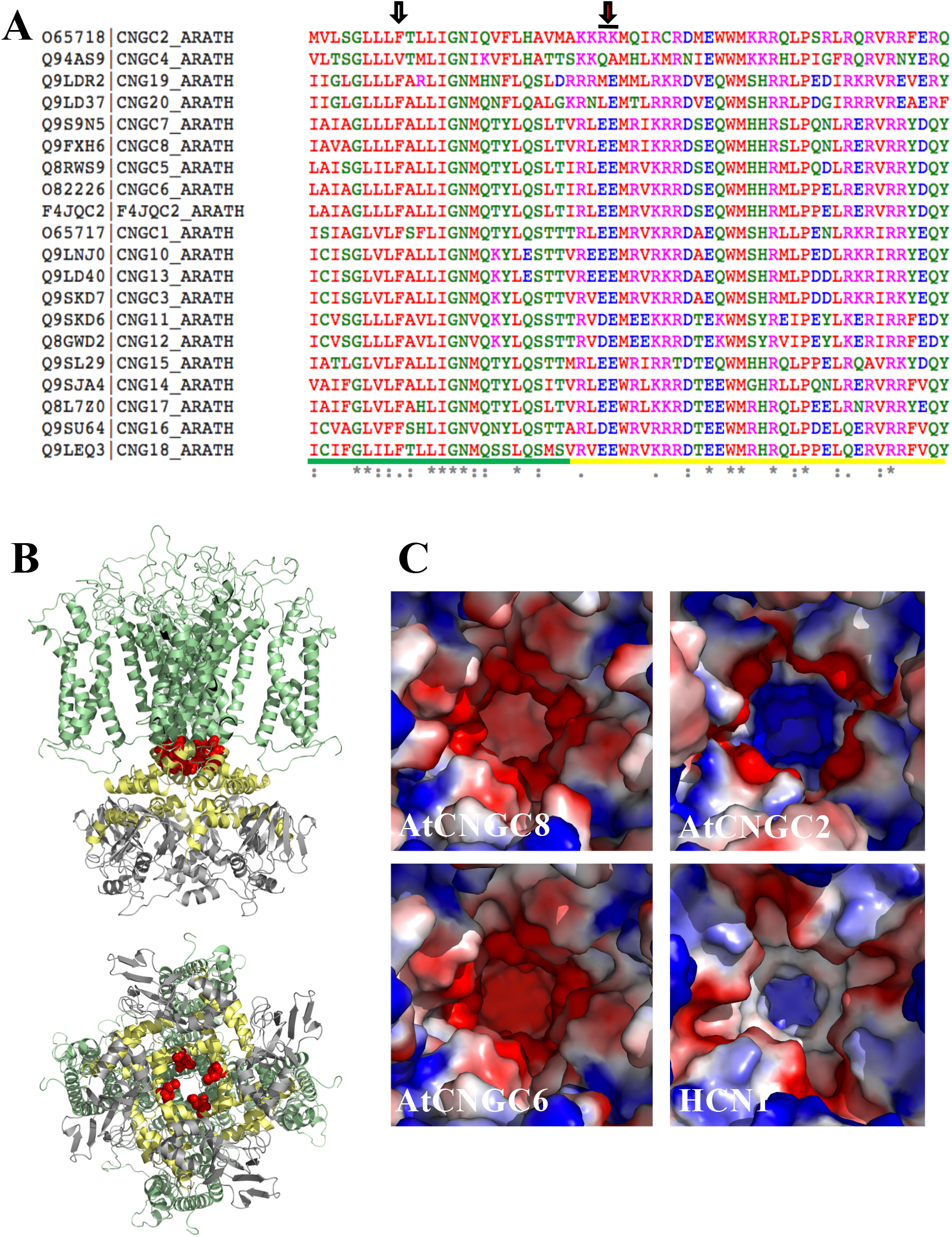
Structural rationale for the channel-blocking effect of Mg^2+^. (**A**) Sequence alignment of the AtCNGC protein sequence surrounding the putative Mg^2+^ binding site. The di-acidic motif present in most AtCNGCs is highlighted by a red arrow and black line, whereas the location corresponding to the Mg^2+^-binding acidic motif in the Kir2.2 pore domain is indicated by the left arrow (grey filling). The residues of the pore and linker regions are underlined green and yellow, respectively. (**B**)Structural model of AtCNGC8, based on human HCN1 (PDB accession number 5u6o). *Top panel*: side view, with the transmembrane region coloured in green, the linker region in yellow, and the cytoplasmic cAMP binding domain in grey. The di-acidic motif (residues E437 and E438 in AtCNGC8) are highlighted as red sphere models. *Bottom panel*: view from the cytoplasm into the channel (90° rotation with respect to top panel). (**C**) Electro-static surface representation (colour-ramped from negatively charged, red, to positively charged, blue) of homology models of AtCNGCs containing the di-acidic motif (AtCNGC8 and 6) and of AtCNGC2 (model) and human HCN1 (PDB 5u6o) lacking this motif.

## DISCUSSION

In order to record ion-currents (for instance *I*_K,in_ or *I*_K,out_) from plant cells in the “whole-cell” patch configuration, it is standard, for reasons highlighted in the introduction (see also: “Methods” section in (50)), to include ATP and Mg^2+^ either in the form of [Mg-ATP] or [MgCl_2_ + K_2_-ATP]. In contrast, the composition of the internal solution used to characterize the hyperpolarization-activated Ca^2+^-current (HACC or *I*_Ba_) is more variable especially with regards to ATP and Mg^2+^. One notices for instance that ATP and Mg^2+^ are either both included (9,11,17,51,52) or completely omitted from the internal solution (10,16,53–55). Here, we carefully addressed the role and consequences of Mg^2+^ inclusion, either in the form of Mg-ATP or MgCl_2_. We report experiments designed to assess the effect of Mg^2+^ on *I*_Ba_ current activated by hyperpolarization in GCPs, being aware that other cell types,, e.g., root cells, might indeed show different responses. We show that omitting Mg-ATP unmasked the presence of an instantaneously-activating inwardly-rectifying conductance. This ‘newly unveiled’ conductance still retains most of the biophysical and pharmacological characteristics that are hallmarks of the classical *I*_Ca_-type (9,10) and CNGCs which we referred to here as either *I*_Ba_ or HACCs (13,14). Like *I*_K,in_, this conductance is activated by hyperpolarizing-going voltages and shows some voltage-dependent rectification (but not as strongly as compared to *I*_K,in_ for instance; see Fig. 4A). Also, typical of HACCs, we found that besides Ba^2+^, other divalent cations such as Ca^2+^, Sr^2+^ and Mn^2+^ also permeate this channel but at a much slower rate than Ba^2+^ does, hence the smaller currents resolved even though using the same amounts of divalent cations in the bath. More importantly, this HACC is activated by cAMP as well as permeable to monovalent cations (see Suppl. Fig. 1 and 2), which are definite attributes of CNGCs. HACCs including CNGCs are specifically blocked by low concentrations of extracellular Gd^3+^ that are far less effective in blocking *I*_K,in_ (see Fig. 4A). Equally important, we found that the unmasked *I*_Ba_ is also enhanced by ≈ 1.3- to 1.5-fold in response to 5 minutes treatment with ABA and up to 2-fold after 10 minutes. Similarly, it was shown that ABA increases a PM *I*_Ca_-type whole-cell current in *Arabidopsis* guard cells by ≈ 2- to 3-fold within 5 minutes of treatment (9). Note that the ABA effect reported here was obtained not only in Mg^2+^-free but even more importantly in ATP-free internal solution, thus indicating that ATP is not as crucial for this channel as was suggested by an earlier report (56). This is, in fact, in agreement with many other reports showing that indeed ABA can increase cytosolic Ca^2+^ levels through activation of PM calcium channels activated by hyperpolarization in *Arabidopsis thaliana* guard cells (10,57,58). Furthermore, ATP was neither required for cGMP-nor cAMP-activated (49,50) Ca^2+^-permeable cation channels in many different plant cell types (mesophyll, guard cells or pollen tubes) that also show many of the HACC characteristics. This observation is in contrast to the above-mentioned report where ATP and subsequent protein (de)-phosphorylation was described as a prerequisite for an ABA effect on calcium channels (56). This indicates that there may be more than one subtype of calcium channels co-existing in the PM and/or that additional modes of regulation of these Ca^2+^-permeable channels are operating which might require ATP- and protein kinases-dependent signaling (56,59).

One discrepancy that stands out in our report is that Mg^2+^ was shown to permeate HACCs in guard cells (10) and in root hairs (51), whereas external Mg^2+^, in our experiments, did not appear to permeate this channel (Fig. 3A). This is even more intriguing, considering the fact that cGMP was recently shown to activate an inward rectifying current (also Lanthanide-sensitive) with Mg^2+^ as a charge carrier (16). Hence this is another hint that we may be dealing with more than one sub-type of calcium channels. In animal cells, Mn^2+^ was described as both a blocker of Ca^2+^-channels, if Ca^2+^ is present in the bath, while in the absence of Ca^2+^, Mn^2+^ permeates the channel (60,61). These data might infer that HACCs, even though sharing many similarities in terms of their biophysical and pharmacological characteristics, might slightly differ from one cell type to another depending on tissue type and/or plant species.

So, the first key finding is that omitting Mg-ATP from the intracellular medium unmasks a larger Gd^3+^-sensitive non-selective cation conductance that is also regulated by cAMP and ABA. The mechanism consists of shifting the I-V characteristic to the right where less negative voltages can mobilize cations (including Ca^2+^) through the channel. The second key finding is the demonstration that Mg^2+^ alone can block this conductance (Fig. 6).

The current activated by voltages more negative than −200 mV is small, but significant (Fig. 1A). At the present time, we have no evidence to support that this instantaneous, rectifying and Mg^2+^-resistant Ba^2+^-current would be carried by a different population type of HACCs (the channel type unmasked when Mg^2+^ was omitted). If anything, this current could still be carried by the same type of channels since addition of 20 μM Gd^3+^ to the bath is still able to swiftly and efficiently block this current.

In summary, our data describe for the first time that in guard cells, Mg^2+^ blocks *I*_Ba_ by shifting the I-V relationship and its activation threshold to more negative voltages (Fig. 1C). This effect is reminiscent of the inhibitory effects by Mg^2+^ on many ion-channels that have been interpreted as ‘charge screening effects’ (18). Indeed, the rectification of the inwardly rectifying K^+^ (Kir) channels is due to a voltage-dependent block by cytosolic Mg^2+^ (and polyamines) thereby blocking outward K^+^-efflux. Upon hyperpolarization, Mg^2+^ is ejected from the pore, which appears to result in a time-dependent opening of the channel (62,63). Likewise, TRPV6 shows Mg^2+^-dependent gating that contributes to its strong inward rectification (20) and it was suggested that Mg^2+^ can block the channel by binding to a site within the transmembrane electrical field where it interacts with other permeant cations (20). It is also conceivable that other mechanisms could be operating such as electrostatic interaction between Mg^2+^ and some PM lipids, such as Phosphatidylinositol 4,5-bisphosphate (or PIP_2_). It was demonstrated that increasing the amount of membrane PIP_2_ results in decreasing the sensitivity of KCNQ channel to inhibition by Mg^2+^ (64). In addition, some Ca^2+^-channels were found to require PIP_2_ for their normal function (65). This begs the question of whether guard cell PM Ca^2+^-channels are also PIP_2_-sensitive. Our results also raise the question of whether Mg^2+^ could be equally important for *I*_K,in_ gating. Indeed, and unlike earlier reports, we found that activation of *I*_K,in_ was dependent on Mg^2+^ being present inside the patch pipet (Fig. 6).

Furthermore, the activation of the above HACCs by hyperpolarization and cAMP, the absence of cation selectivity and inhibition by lanthanides is consistent with the hypothesis that channels responsible for the observed effects are CNGCs and this is consistent with our structural modelling that has revealed the presence of di-acidic motifs in the pore forming helix of a subset of AtCNGCs. Acidic residues pointing towards the inner side of the pore, have previously been shown to confer Mg^2+^-dependence to inward rectifying K^+^ channels in animals (47) (Suppl. Fig. 3A). Given the position of the di-acidic motif in the cytoplasmic side of the pore, it is conceivable that Mg^2+^-binding can be affected by changes in pore opening, for example introduced by cAMP binding to the cytoplasmic region of AtCNGCs. Such a crosstalk would provide a mechanistic explanation for our observation that db_2_cAMP overrides the channel blockage produced by 1mM MgCl_2_ (no added ATP) (Suppl. Fig. 2B).

We therefore propose, firstly, that Mg^2+^ can limit, if not prevent, continuous Ca^2+^-leakage possibly through all HACCs (including CNGCs) at resting membrane potentials and secondly, that the activation of these channels requires mechanism(s) by which Mg^2+^-binding is altered, such in the case of adding cAMP (Suppl. Fig. 2B), in turn giving Mg^2+^ an important role in calcium homeostasis and calcium-dependent signaling.

## Acknowledgements

This research has been supported by the King Abdullah University of Science and Technology (KAUST). We are indebted to Professor Enid MacRobbie (Department of Plant Science, University of Cambridge, UK) for allowing us to use some of the data gathered by FL-C while in her laboratory (research was supported by BBSRC Grant P05730 to E.M.). We also thank Prof. Mark Tester for his invaluable comments.

## Author Contributions

FL-C and CG conceived the study, FL-C performed the experiments and analyzed the data. CG and SA performed the structural analyses. All authors contributed to the writing of the manuscript.

## Competing financial interests

The authors declare no competing financial interests.

---

**Suppl. Figure 1.**
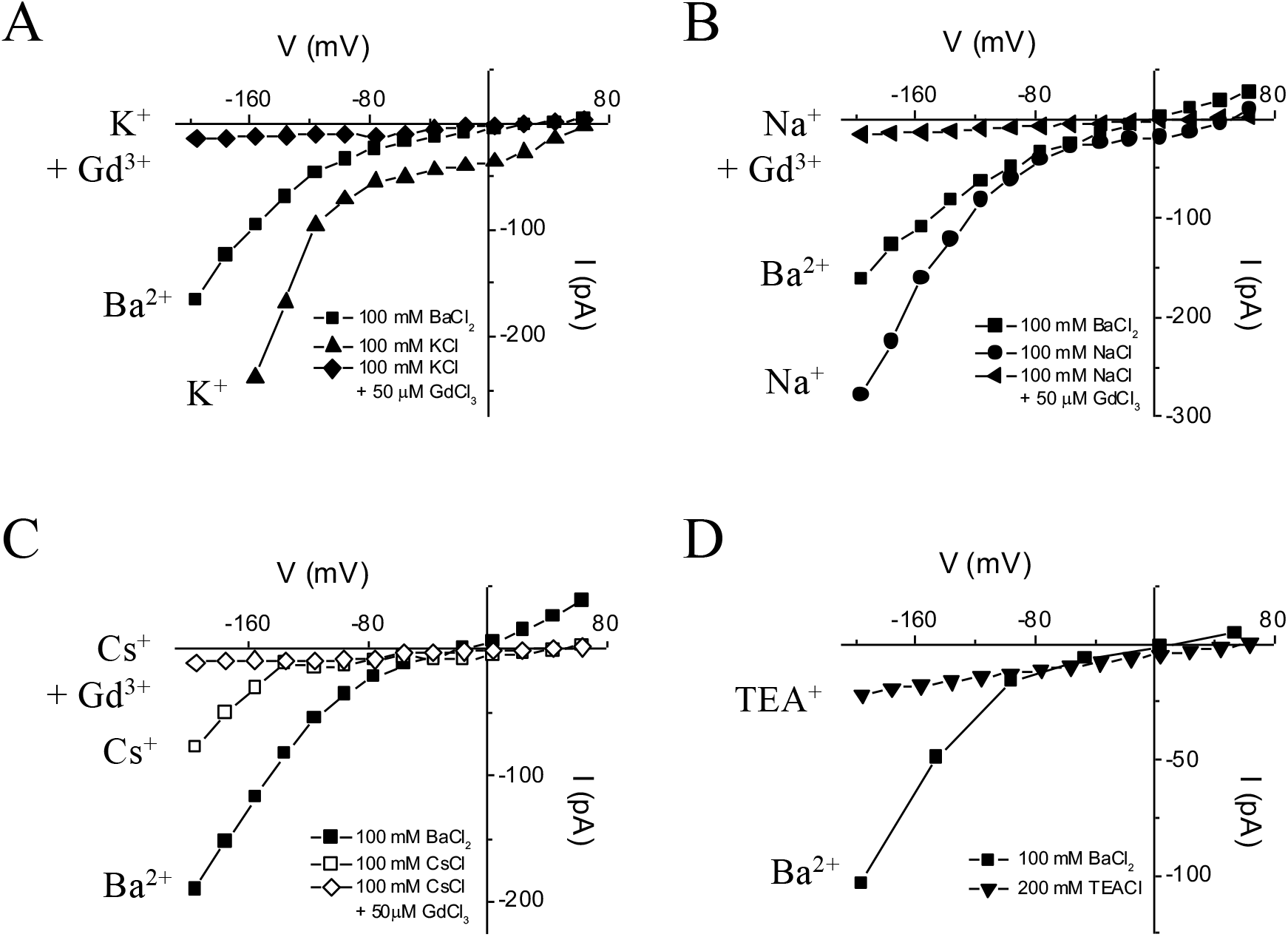
In the absence of MgATP, Guard cell’s HACCs are permeable to monovalent cations such as K^+^, Na^+^, Cs^+^ but not TEA^+^. All experiments were conducted in the whole cell configuration where *V. faba* GCPs were held at −56 mV. (**A**) Superimposed I-V plots in the presence of 100 mM BaCl_2_ (■), 100 mM KCl (▲) or 100 KCl + 0.05 mM GdCl_3_. (**B**) Superimposed I-V plots in the presence of 100 mM BaCl_2_ (■), 100 mM NaCl (⚫) or 100 NaCl + 0.05 mM GdCl_3_. (**C**) Superimposed I-V plots in the presence of 100 mM BaCl_2_ (■) or 100 mM CsCl (□) or 100 CsCl + 0.05 mM GdCl_3_ (◊). (**D**) Superimposed I-V plots in the presence of 100 mM BaCl_2_ (■) or 100 TEACl (▼).

**Suppl. Figure 2.**
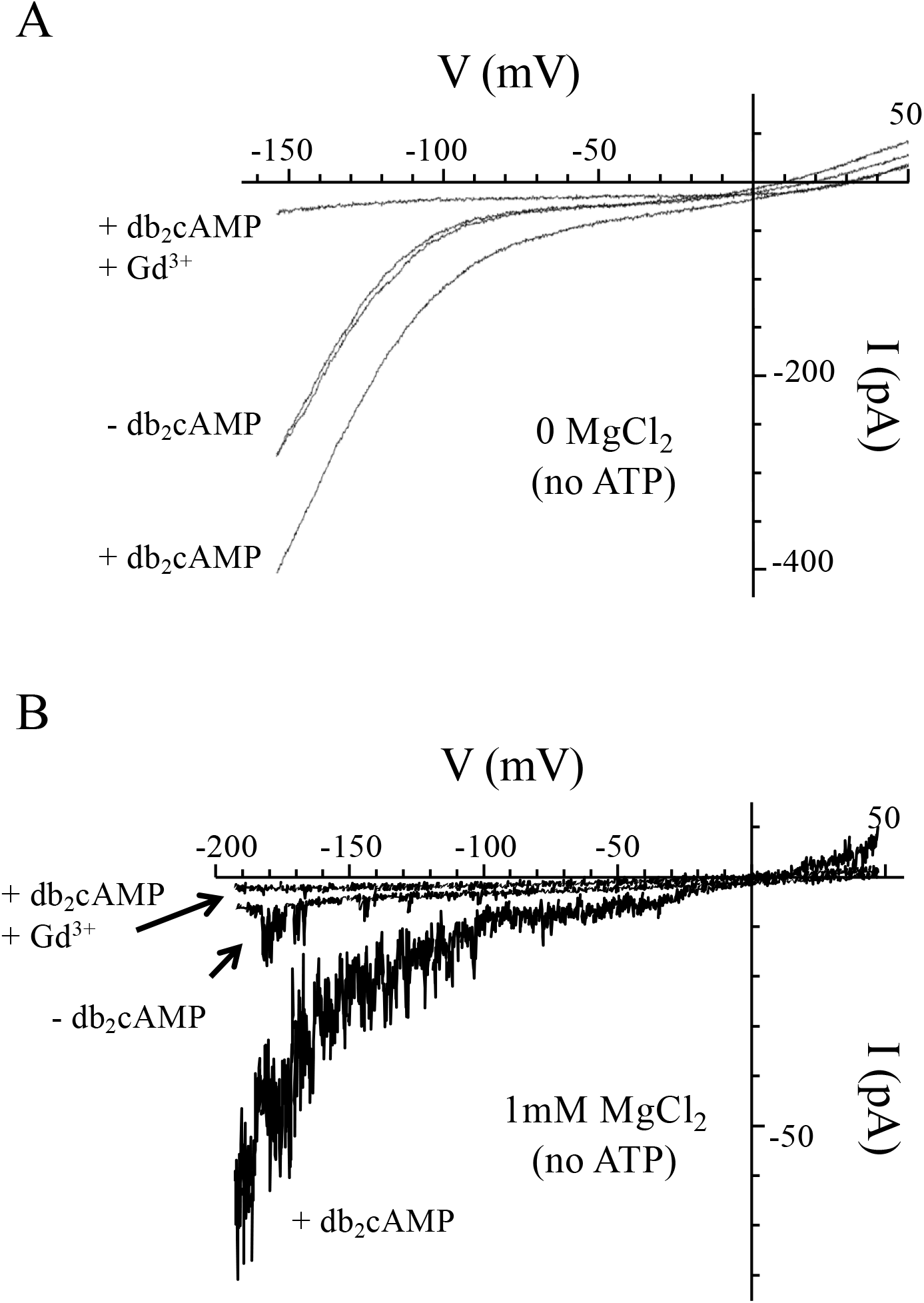
Dibutyryl cyclic AMP potentiates a Gd^3+^-sensitive current in guard cells either in the presence or absence of intracellular Mg^2+^. Experiments were conducted in the whole cell configuration where the GCPs were held at −52 mV and the I-V plots were generated using a hyperpolarizing ramp protocol. see Methods for bath and intracellular media. (**A**) Typical example from a *V. faba* GCP patched with no MgATP in the intracellular media showing superimposed I-V ramps from +50 to −156 mV (70 mV.s^−1^) in the absence (-db_2_cAMP) or presence (+ db_2_cAMP) of 1 mM dibutyryl cyclic AMP. Note that adding 0.05 mM GdCl_3_ while keeping db_2_cAMP in the bath blocks this conductance. (**B**) Typical example from an *A. thaliana* GCP patched with 1mM MgCl_2_ (no added ATP) in the intracellular media showing superimposed I-V ramps from ~50 to −192 mV (70 mV.s^−1^) in the absence (−db_2_cAMP) or presence (+ db_2_cAMP) of 1 mM dibutyryl cyclic AMP. Note that adding 0.05 mM GdCl_3_ while keeping db_2_cAMP in the bath also blocks this conductance.

**Suppl. Figure 3.**
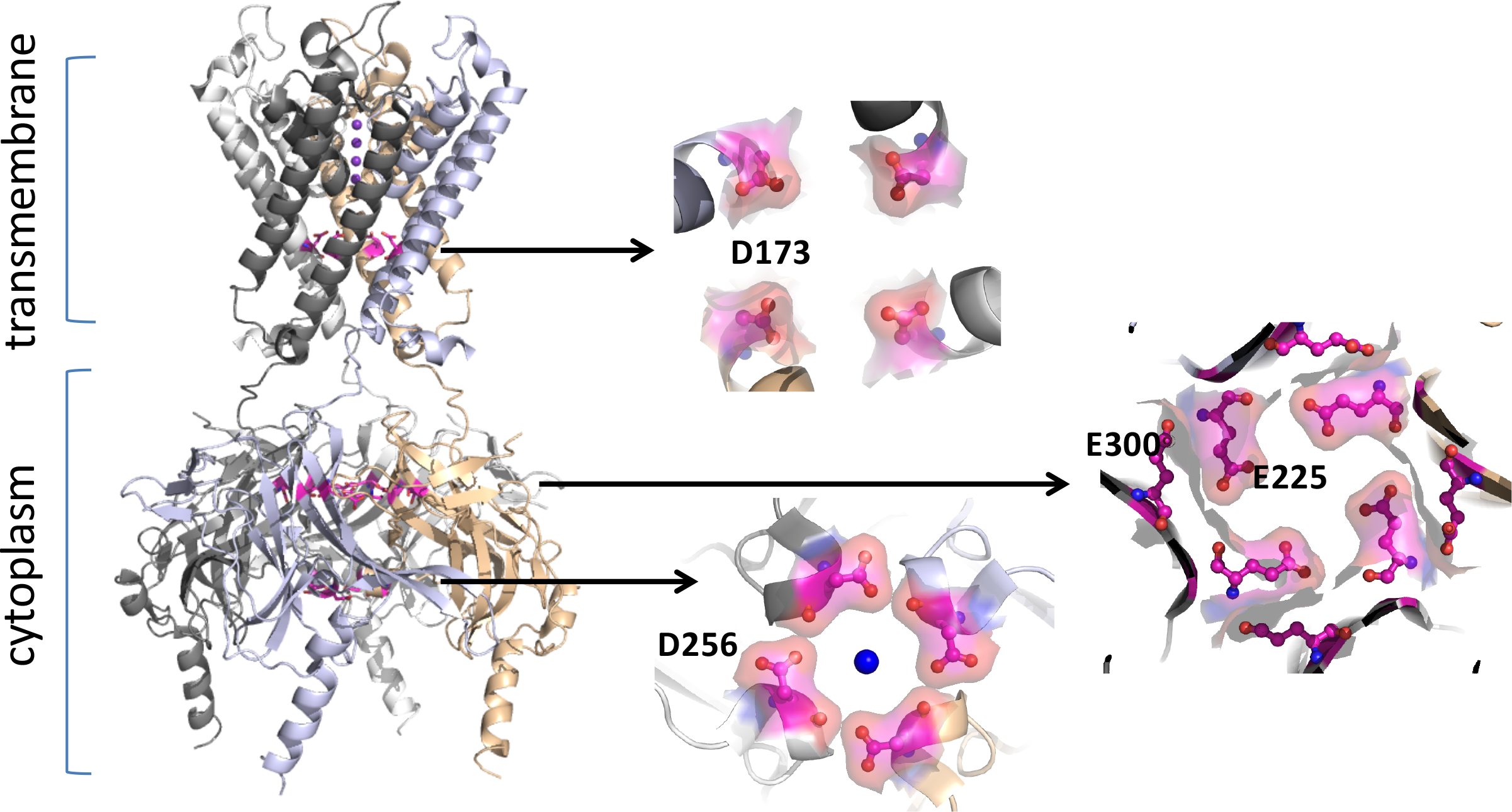
Mechanism of inward rectifying by magnesium ions. Tao *et al*. (Science 2009, 326, 1668−1674) have shown that inward rectifying through Mg^2+^ can be explained by the ion binding to negatively charged regions in the pore (formed by D173) and in the cytoplasmic regulatory domains (D256 and E300/E225). The crystal structure of the inward rectifying potassium channel Kir2.2 (Tao, 2009; PDB entry 3JYC) is shown in ribbon presentation. The four subunits are colour-coded. Potassium ions in the channel are shown as magenta spheres. Negatively charged residues that bind the Mg^2+^ - mimic Sr^2+^ in the crystal structure are shown in pink in their molecular surface.

